# Toward Single-Cell Control: Noise-Robust Perfect Adaptation in Biomolecular Systems

**DOI:** 10.1101/2025.10.20.683581

**Authors:** Dongju Lim, Seokhwan Moon, Yun Min Song, Minjun Kim, Jinyeong Kim, Kangsan Kim, Byung-Kwan Cho, Jinsu Kim, Jae Kyoung Kim

## Abstract

Robust perfect adaptation (RPA), whereby a consistent output level is maintained even after a disturbance, is a highly desired feature in biological systems. This property can be achieved at the population average level by combining the well-known antithetic integral feedback (AIF) loop into the target network. However, the AIF controller amplifies the noise of the output level, disrupting the single-cell level regulation of the system output and compromising the conceptual goal of stable output level control. To address this, we introduce a new regulation motif, the noise controller, which is inspired by the AIF loop but differs by sensing the output levels through the dimerization of output species. Combining this noise controller with the AIF controller successfully maintained system output noise as well as mean at their original level, even after the perturbation, thereby achieving noise RPA. Furthermore, our noise controller could reduce the output noise to a desired target value, achieving a Fano factor as small as 1, the commonly recognized lower bound of intrinsic noise in biological systems. Notably, our controller remains effective as long as the combined system is ergodic, making it applicable to a broad range of networks. We demonstrate its utility by combining the noise controller with the DNA repair system of *Escherichia coli*, which reduced the proportion of cells failing to initiate the DNA damage response. These findings enhance the precision of existing biological controllers, marking a key step toward achieving single-cell level regulation.

## Introduction

All biological systems experience fluctuations and perturbations, such as changes in protein levels induced by differing temperatures^1^. Maintaining a stable output despite such perturbations is desirable across diverse living systems^2^ and the process is known as robust perfect adaptation (RPA)^3-5^. Achieving RPA through engineered biological modules has been a focus of many studies^6-10^. Among these, one of the most promising modules is the antithetic integral feedback (AIF) loop^6^. Integrating an AIF module allows a system to maintain stable output levels even after perturbations, independent of reaction parameter values.

However, the AIF cannot be applied to single-cell level controllers as it regulates the system output only at an “average” level. For instance, if the AIF controller regulates protein P at a mean level of 10 across 100 cells, the population average of P over 100 cells remains at 10. However, individual cells may exhibit significant variation: one cell might have P = 1 while another cell has P = 20. This intrinsic noise poses a challenge for single-cell control, which is critical in biological systems. For example, in cancer therapy, even if 99% of cancer cells are eliminated, the remaining 1% can proliferate, reducing treatment efficacy^11^. Similarly, tuberculosis prevention requires eliminating all *Mycobacterium tuberculosis* cells, as even a few survivors can lead to disease reactivation^12^. Addressing these challenges requires not only maintaining the population-level mean but also controlling noise at the single-cell level^13^. However, the AIF controller cannot regulate noise; instead, it amplifies the noise compared to the original system^14,15^. As a result, while the AIF controller preserves the population average, the increased noise limits its practical utility for single-cell control.

To overcome this limitation, previous studies have modified the AIF controller or incorporated additional modules to suppress noise^14-16^. For example, Briat et al. and Filo et al. effectively reduced the noise by incorporating additional negative feedback loops^14,16^, while Kell et al. directly modified the AIF controller to achieve noise reduction^15^. However, no module has yet been developed that is capable of maintaining what we call ‘noise RPA’, which is maintaining the output noise level at its original level even after the perturbation in the system.

In this study, we developed a new controller module, the noise controller (NC), which enables noise RPA. By combining both the AIF controller and NC into a system, we achieved RPA for both mean and noise, maintaining the output mean and noise at its original level even after the perturbation. Beyond maintaining the output noise, we further reduced the noise up to a Fano factor of 1. This capability is preserved as long as the combined system is ergodic with finite second moments, making it applicable across diverse biological networks. Furthermore, we illustrate how combining the AIF controller and NC can reduce the failure cases in the DNA damage response. By suppressing system noise, our NC significantly enhances the precision of biological controllers. Therefore, NC provides a powerful strategy for achieving single-cell level control, which has been unattainable with the AIF controller alone.

## Results

### The noise controller achieves noise robust perfect adaptation

The AIF controller is the robust mean controller (MC)^6,7^ (Fig. 1a). By combining this MC with the original network without the RPA property, the mean of the output level (i.e., X_2_ in Fig. 1a) can be maintained even after a perturbation in the system, thereby achieving RPA for the mean (mean RPA) (Fig. 1b). However, this approach does not guarantee RPA for the noise (noise RPA), and amplifies the noise levels (Fig. 1c). Due to this noise amplification, the output level remains inconsistent despite the MC’s regulation of the mean output level, undermining the practical significance of the mean RPA. To address this problem, a new module is needed to preserve the original noise level.

**Figure 1.**
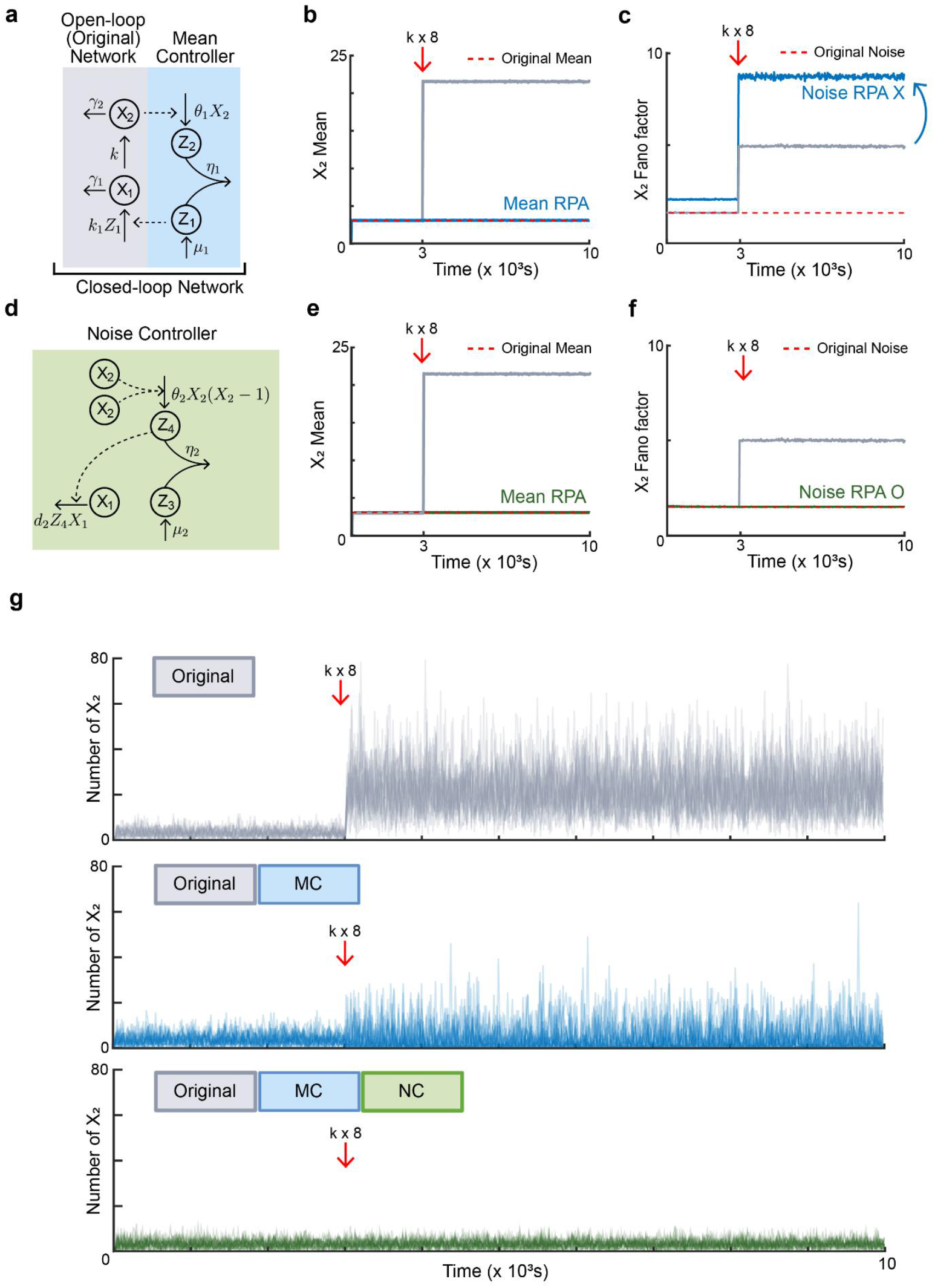
The noise controller achieves noise RPA. **(a)** The mean controller (MC) can be integrated into the original open-loop network to form a closed-loop network. **(b-c)** The mean (b) and noise (Fano factor) (c) of 50,000 trajectories of *X*_2_ (output) from the original network (gray) and original network with MC (blue) simulated with parameters *γ*_1_ = 1, *γ*_2_ = 3, *k* = 2, *μ*_1_ = 3, *θ*_1_ = 1, *η*_1_ = 50, *k*_1_ = 1 and *k* increased 8-fold at time = 3000s. The original network with MC achieves robust perfect adaptation (RPA) for the mean (b, blue), maintaining its original mean output level (b, red dotted line) even after a parameter perturbation (b, red arrow), in contrast to the original network alone (b, gray). However, the output noise (i.e., Fano factor of *X*_2_) of the original network with MC (c, blue) does not exhibit RPA, and increases compared to the original network both before and after the perturbation (c, gray). **(d)** To maintain both the output noise and the mean of the output level at the original levels, also after a perturbation, we propose the noise controller (NC). NC differs from MC in that it senses the second moment of *X*_2_ by the dimerization of *X*_2_ (the reaction labelled *θ*_2_) and promotes the degradation of *X*_1_ (See Methods for more details). **(e-f)** The mean (e) and Fano factor (f) of 50,000 trajectories of *X*_2_ from the original network (gray) and original network with MC and NC (green) simulated with parameters following those in Fig. 1b-c and additional parameters, *μ*_2_ = 9.05, *θ*_2_ = 1, *η*_2_ = 50, *d*_2_ = 1. The original network combined with MC and NC achieves RPA for not only the mean output level (e, green line) but also the output noise levels (f. noise RPA). **(g)** Overlay of five single-cell trajectories of the copy number of *X*_2_ from the original network (gray, top), original network with MC (blue, middle), and original network with MC and NC (green, bottom) simulated with parameters following those in Fig. 1e-f. Unlike the original network and the original network with MC, the original network with MC and NC achieves RPA for both mean and noise levels. All the simulation results are obtained using the Gillespie algorithm and the Monte Carlo method.

To achieve noise RPA as well as mean RPA, we developed a new module, the noise controller (NC) (Fig. 1d). NC is inspired by MC but is designed to sense the second moment of the output level (the mean of the squared number of output species, 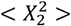), whereas MC senses the first moment (*X*_2_). Therefore, using both MC and NC will enable control over the output noise level (i.e., Fano factor, 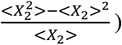 (See Methods for more details).

To test this, we simulated the output noise level of the original network combined with MC and NC before and after the perturbation in the system. A simulation result shows that combining both MC and NC preserves both the mean (Fig. 1e) and noise (Fig. 1f) at its original level, across a wide range of perturbation magnitudes (Fig. S4). Therefore, by combining MC and NC, both mean RPA and noise RPA can be achieved, in contrast to the MC alone, which only achieved mean RPA (Fig. 1g).

### The noise controller reduces noise up to Fano factor 1

While combining the MC with the original network amplifies the output noise, we successfully recovered this noise to its original level by further combining NC (Fig. 2a). This is possible because MC controls the first moment and NC controls the second moment, ultimately controlling the noise (Fano factor). Here, the value of the Fano factor, 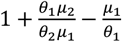, is determined by the reaction parameter of MC and NC (i.e., *θ*_1_, *μ*_1_, *θ*_2_ and *μ*_2_ in Fig. 1a and d; See Methods for more details). Given this, adjusting the parameter values of MC and NC may allow us to reduce the noise beyond the original level as much as desired (Fig. 2a).

**Figure 2.**
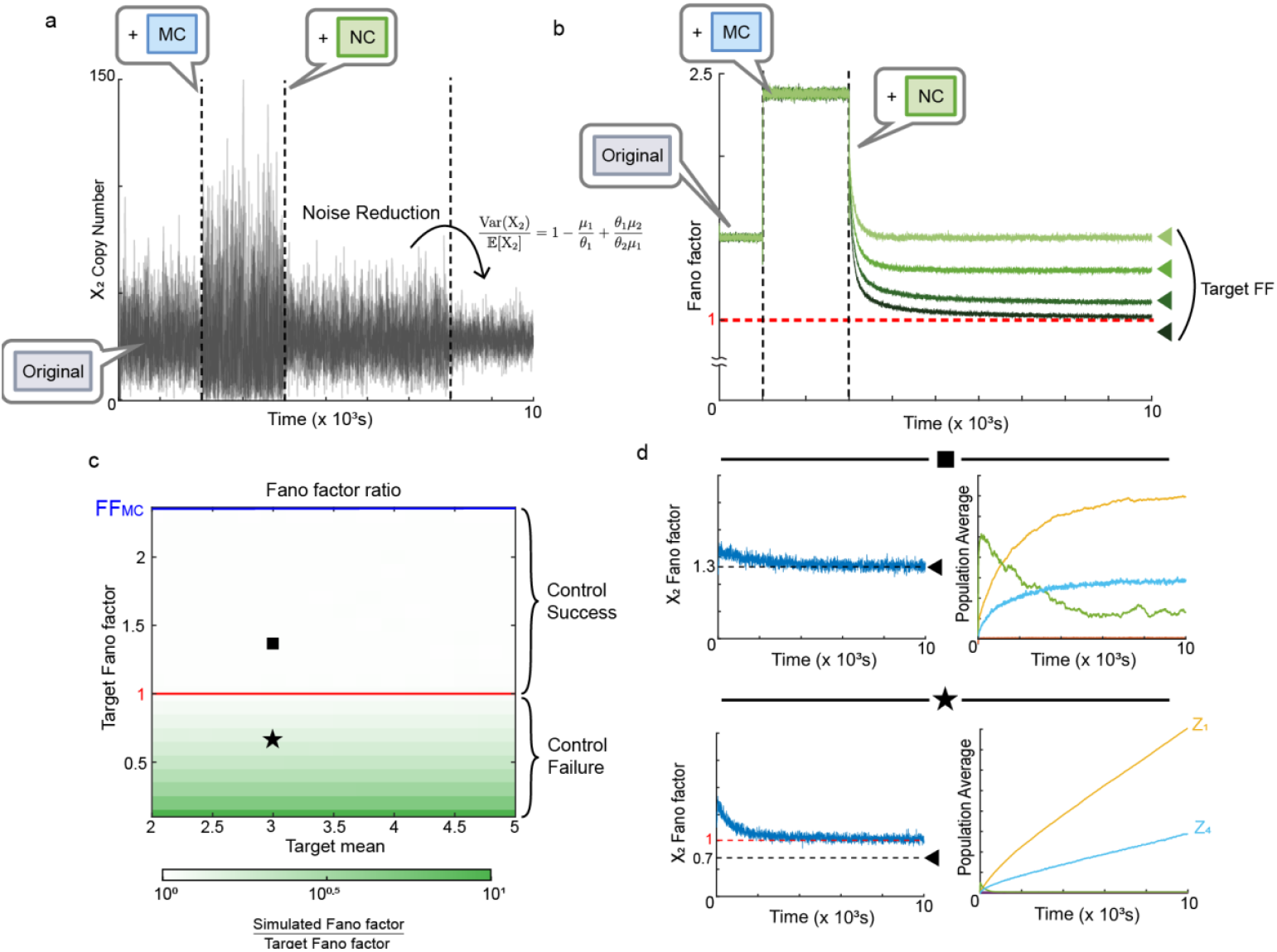
The noise controller can reduce output noise below the original level, up to a Fano factor of 1. (a) Twenty trajectories of *X*_2_ (output) from the original network (time = 0 – 2000s), original network with MC (time = 2000 – 4000s), and original network with MC and NC (time = 4000s – 8000s) simulated with parameters following those in Fig. 1e-f except for *k* = 20, *μ*_1_ = 27, *μ*_2_ = 829.05 and *μ*_2_ was decreased to *μ*_2_ = 729.05 at time = 8000s. Combining the MC with the original network led to an increase in output noise, which was restored to the level of the original network by further combining the NC. The output noise could be reduced below its original level by adjusting the key parameters in the MC (*θ*_1_ and *μ*_1_ in Fig. 1a) and NC (*θ*_2_ and *μ*_2_ in Fig. 1d) since the expected Fano factor with NC is 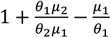. **(b)** The Fano factor of 50,000 trajectories of *X*_2_ from the original network (time = 0 – 1000s), original network with MC (time = 1000 – 3000s), and original network with MC and NC (time =3000s –) simulated with parameters following those in Fig. 1e-f, while *μ*_2_ was adjusted to achieve the target Fano factor. When one of the key parameters in NC, *μ*_2_, was adjusted to 8.7, 9.3, 9.9, 10.5 to achieve target Fano factor 2.7/3, 3.3/3, 3.9/3, 4.5/3, respectively, NC successfully reduced the output noise to the target Fano factors (triangles) for the target values greater than 1 (top three triangles). However, for the target Fano factor below 1 (the lowest triangle), the output noise failed to reach the target and stabilized at the Fano factor of 1 (red dotted line). **(c)** A heatmap visualizing the ratio of simulated output noise from the original network with MC and NC (simulated Fano factor) to the target output noise (target Fano factor) across different values of target value for output mean (target mean) and target Fano factor. These values were adjusted by modifying key parameters in MC and NC, *θ*_1_ in Fig. 1a and *θ*_2_ in Fig. 1d, while remaining parameters were consistent with Fig. 1e-f. Here, FF_MC_ represents the Fano factor of the original network with MC. The heatmap demonstrates that the NC successfully achieves the target when the target Fano factor is greater than 1 (the area above the red line). **(d)** For target Fano factors greater than 1 (square), the output noise converges to the target value (black dotted line; left) and other species in the system reach steady states (right). In contrast, for target Fano factors below 1 (star), the output noise stabilizes at the Fano factor of 1 (red dotted line) and fails to achieve the target value (black dotted line; left). This is because certain species in the system, particularly the *Z*_1_ and *Z*_4_, diverge and fail to control the system effectively (right). All the simulation results are obtained by using the Gillespie algorithm and the Monte Carlo method.

To investigate whether this noise reduction is achievable, we adjusted the parameters of MC and NC so that the mean output level remained fixed at the original network level, while the Fano factor was set below the original noise level (Fig. 2b). The results showed that when the target Fano factor was greater than 1, the output noise was successfully reduced to the target value. In contrast, when the target Fano factor was smaller than 1, the output noise did not decrease below the Fano factor of 1, and the desired noise reduction was not achieved. This suggests that MC and NC allow us to reduce the output noise as much as desired up to a Fano factor of 1.

We further examined whether this ability of MC and NC holds even when the target mean output level differs from the original level (Fig. 2c). To do this, we adjusted MC and NC for varying target means and target Fano factors, combined them with the original network, and simulated the Fano factor of this combined network. Then, the ratio between the simulated Fano factor and the target Fano factor was computed and visualized as a heatmap (Fig. 2c). The results showed that regardless of the target mean, when the target Fano factor was greater than 1, the simulated Fano factor was identical with the target Fano factor. This indicates that the output noise can be reduced as much as desired if the target Fano factor is greater than 1. In contrast, when the target Fano factor is smaller than 1, the simulated Fano factor remained higher than the target Fano factor, meaning the desired noise reduction was not achieved. This occurs because some controller species (Z_1_ in Fig. 1a and Z_4_ in Fig. 1d) diverged when the target Fano factor was set below 1 (Fig. 2d), failing to function properly as controllers in contrast to the case when the target Fano factor was greater than 1 (Fig. 2d). A detailed explanation of why Z_1_ and Z_4_ diverge is provided in Fig. S3 and Text S2.

### Noise reduction can be achieved for various networks over a broad range of parameter conditions

We have shown that the output noise can be reduced down to a Fano factor of 1 by adjusting key reaction parameters of MC and NC (Fig. 2). In addition to these key parameters, MC and NC also have parameters related to the sequestration reaction (*η*_1_ and *η*_2_) and actuation reaction (*k*_1_ and *d*_2_). Thus, we investigated how variations in these parameters affect the noise reduction capability of MC and NC (Fig. 3a,b).

**Figure 3.**
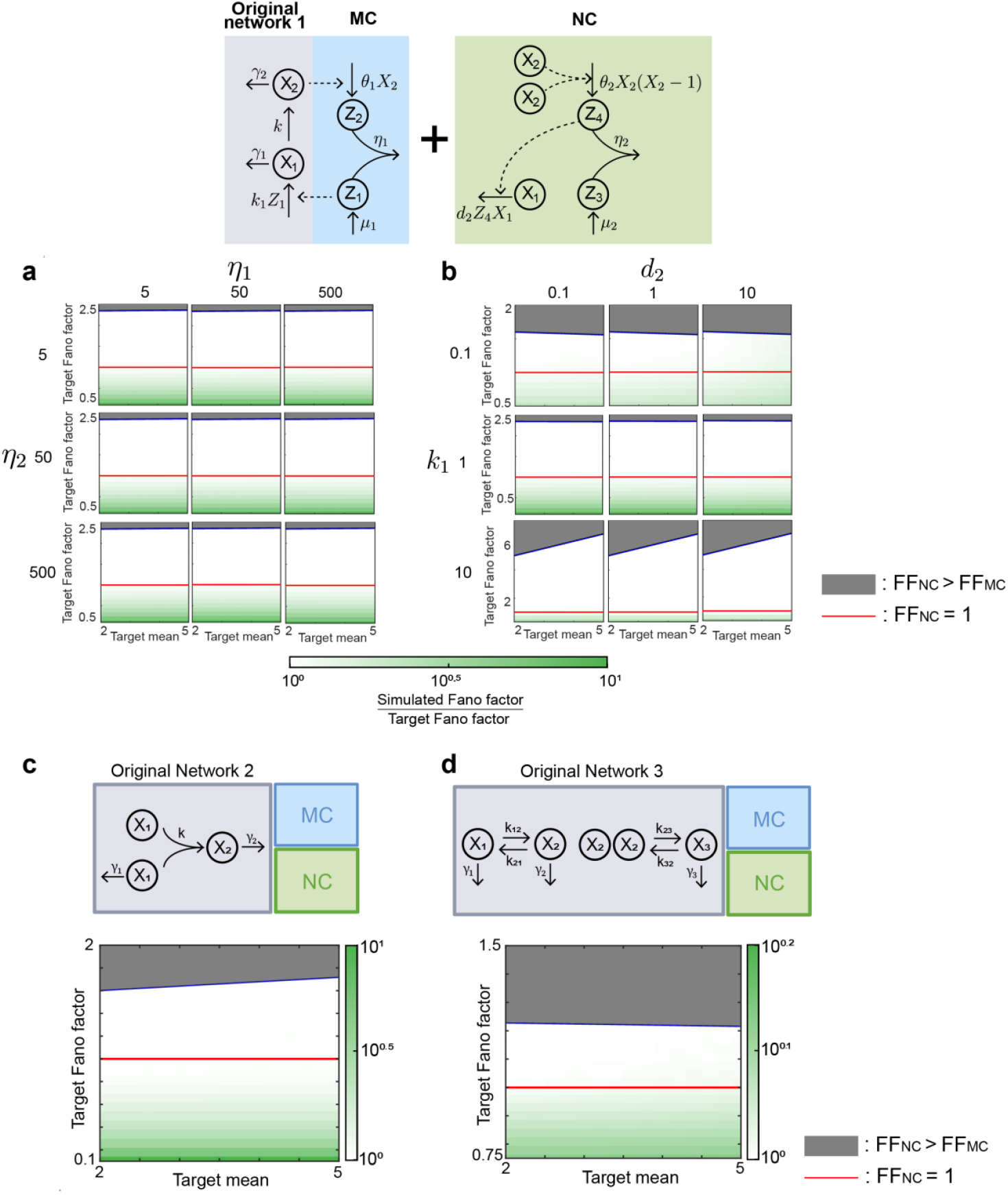
The NC can reduce the output noise up to a Fano factor of 1 across various parameter values and original network structures. **(a-b)** Heatmaps visualizing simulations from Fig. 2c with varying sequestration parameters, *η*_1_ and *η*_2_, (a) and actuation parameters, *k*_1_and *d*_2_ (b). Each parameter was increased or decreased 10-fold from the original value (*η*_1_ = *η*_2_ = 50, *k*_1_ = *d*_2_ = 1). The gray shaded region indicates that the target Fano factor (FF_NC_) is greater than the Fano factor of the original network with MC (FF_MC_). The analysis focuses on the area where the FF_NC_ is less than the FF_MC_ (i.e., the area outside the gray-shaded region), in agreement with the objective of noise reduction. Regardless of the values of the sequestration and actuation parameters, NC successfully reduces the output noise up to a Fano factor of 1. **(c-d)** A heatmap visualizing simulations from Fig. 2c using alternative networks, which involve bimolecular reactions (see Methods for details). NC successfully reduces the output noise up to a Fano factor of 1. See Methods for more details on stochastic simulations.

To test this, we simulated the heatmap in Fig. 2c using various combination of sequestration parameters (Fig. 3a) and actuation parameters (Fig. 3b). When we varied the sequestration parameters of MC and NC (*η*_1_ and *η*_2_) from the original values of *η*_1_ = *η*_2_ = 50 to 0.1-fold and 10-fold, covering nine different cases, we successfully reduced the noise up to a Fano factor of 1 in all 9 cases (Fig. 3a). Similarly, when the actuation parameters (*k*_1_ and *d*_2_) were varied from the original values of *k*_1_ = *d*_2_ = 1 to 0.1-fold and 10-fold, noise reduction to a Fano factor of 1 was still achievable in all cases (Fig. 3b). These results demonstrate that MC and NC can reduce the output noise of the original network up to a Fano factor of 1, regardless of the sequestration and actuation parameters.

Additionally, we investigated whether MC and NC can reduce output noise in different original networks (Fig. 3c,d). To do this, we simulated the heatmap in Fig. 2c for two new networks: original network 2, which represents a dimerization process (Fig. 3c), and original network 3, which includes a bimolecular reaction process (Fig. 3d). The results showed that we can reduce the noise of both original networks up to a Fano factor of 1.

In conclusion, the noise reduction capability of MC and NC holds across various parameter conditions and various original networks. This is because the Fano factor of the total system (original network with MC and NC) converges to a value that solely depends on the key reaction parameters of MC and NC (Fig. 2a), provided that the total system is ergodic with finite second moments (see Methods for details).

### MC and NC prevent bimodality in output distribution

Using MC and NC, the output noise of the original network can be reduced up to a Fano factor of 1. However, a reduced Fano factor value alone cannot guarantee our ultimate goal of single-cell control. For example, if the output distribution is bimodal, then the output level at the single-cell level may deviate from the desired set point even with low output noise. Therefore, we investigated whether MC and NC can also prevent bimodality in output distribution (Fig. 4).

**Figure 4.**
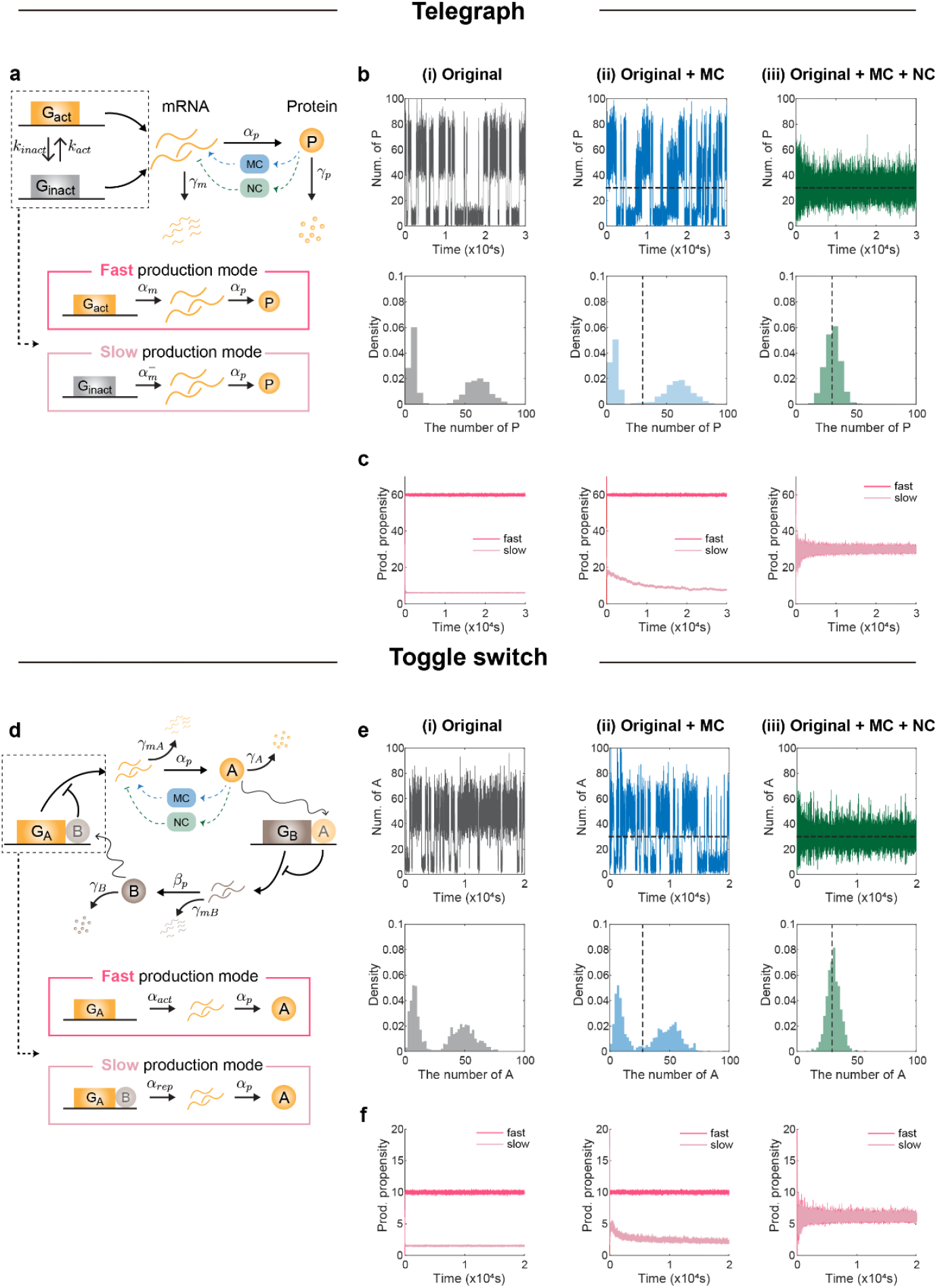
NC prevents bimodality in the output stationary distribution. **(a)** Schematic diagram of the telegraph model with MC (blue) and NC (green). The model alternates between two protein (*P*) production modes (dotted box): a fast production mode (pink box), in which an active gene (*G*_*act*_) produces mRNA and subsequently protein, and a slow production mode (pale pink box), in which an inactive gene (*G*_*inact*_) produces mRNA and subsequently protein at a lower rate. **(b)** A representative single-cell trajectory of *P* (top) and the histogram of its stationary distribution (bottom) simulated from the telegraph model (i), telegraph model with MC (ii), and the telegraph model with MC and NC (iii). Simulations were performed with parameters *k*_*act*_ = 10^−3^, *k*_*inact*_ = 10^−3^, *α*_*m*_ = 60, 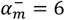, *α*_*p*_= 1, *γ*_*m*_= 1, *γ*_*p*_= 1 and parameters of MC and NC following Fig. 1 except *μ*_1_ =30, *μ*_2_ = 906. See Methods for more details on stochastic simulations. In the original telegraph model (i) and with MC alone (ii), *P* switches between high (∼ 60) and low (∼ 6) levels (i and ii, top), producing a bimodal stationary distribution (i and ii, bottom), whereas with both MC and NC (iii), *P* no longer switches between two distinct states and it shows decreased deviation from the target mean (dotted lines) value (iii, top), resulting in a unimodal stationary distribution (iii, bottom). **(c)** The average production propensity of *P* when the gene is active (pink) or inactive (pale pink), calculated over 1,000 trajectories simulated from the telegraph model (left), the telegraph model with MC (center), and the telegraph model with MC and NC (right). The difference between fast and slow production propensities is large for the telegraph model (left) and the telegraph model with MC (center), but this difference is eliminated when NC is additionally included (right). **(d)** Schematic diagram of the toggle switch model with MC (blue) and NC (green). The model alternates between two protein production modes (dotted box): a fast production mode (pink box), in which an active gene (*G*_*A*_) produces mRNA and subsequently protein (*A*) at a higher rate, and a slow production mode (pale pink box), in which the repressed gene (*G*_*A*_ bind with *B*) produces mRNA and subsequently *A* at a lower rate. **(e-f)** The same analyses as (b–c), but with the telegraph model replaced by the toggle switch model. Simulations were performed with parameters *α*_*act*_ = 10, *α*_*rep*_ = 1.5, *α*_*p*_ = 1, *γ*_*mA*_ = 1, *γ*_*A*_ = 0.2, *β*_*p*_ = 1, *γ*_*mB*_ = 1, *γ*_*B*_ = 0.1 and parameters of MC and NC following Fig. 1 except *μ*_1_ = 30, *μ*_2_ = 906. See Methods for more details on stochastic simulations. The results are consistent with those obtained from the telegraph model.

To do this, we combined MC and NC with a telegraph model^17^, which exhibits a bimodal stationary distribution in its output, and observed how the stationary distribution of the output changed (Fig. 4a– c). In the telegraph model, protein synthesis occurs in two modes: the fast production mode, where active gene produces protein rapidly, and the slow production mode, where the inactive gene produces protein slowly (Fig. 4a). The two modes alternately appear at the single-cell level, causing the protein amount to fluctuate between high and low states (Fig. 4b (i), top), leading to a bimodal stationary distribution (Fig. 4b (i), bottom). Even when MC was combined to control the target mean, protein levels at the single-cell level still alternated between high and low states with a large deviation from the target (Fig. 4b (ii), top), and the stationary distribution remained bimodal (Fig. 4b (ii), bottom). In contrast, when both MC and NC were combined, protein levels fluctuated only around the desired set point without sharp transitions (Fig. 4b (iii), top), and the stationary distribution became unimodal (Fig. 4b (iii), bottom).

To understand why the stationary distribution became unimodal when MC and NC were combined, we investigated how the protein production propensity (i.e., reaction *α*_p_ in Fig. 4a) in the fast and slow production modes changed with the addition of MC and NC (Fig. 4c). In particular, the propensity for the fast production mode was calculated from 1,000 simulated protein production propensity trajectories by averaging, at each time point, the values corresponding to instances in which the gene was active. Applying the same method to cases where the gene was inactive, we obtained the propensity in the slow production mode (see Methods for details). In the original telegraph model, the difference between the propensities of the fast and slow modes was pronounced (Fig. 4c, left), and this difference persisted even when MC was combined (Fig. 4c, center). However, when both MC and NC were combined, the fast-mode propensity decreased, while the slow-mode propensity increased, eliminating the gap between the two modes (Fig. 4c, right).

The convergence of the two production modes by MC and NC arises from the simultaneous action of production-based actuation from MC and degradation-based actuation from NC (see Fig. S1 and Discussion for details). Production-based actuation increases mRNA, raising the slow-mode propensity, while degradation-based actuation reduces mRNA, directly lowering the fast-mode propensity. As MC alone lacks degradation-based actuation, without NC, the fast-mode propensity can only decrease through natural mRNA degradation. As a result, the fast-mode propensity does not sufficiently decrease in the original telegraph model or with MC alone (Fig. 4c (i-ii)), which in turn constrains any increase in slow-mode propensity, as such an increase would otherwise raise the output mean beyond the target value. Thus, a large difference remains between the propensities of the two modes in the original telegraph model or with MC alone (Fig. 4c (i-ii)), leading to large protein-level differences during mode switching. In contrast, when both MC and NC are used, the two propensities become nearly identical (Fig. 4c (iii)), so protein levels show little difference during mode switching.

This pattern was also observed when MC and NC were combined with the toggle switch model^18^ (Fig. 4d–f)—another model that exhibits a bimodal stationary distribution due to fast and slow production modes. Taken together, for the systems with two output production modes, combining both MC and NC can prevent bimodality by making the two modes of the bimodal distribution identical.

### MC and NC reduce the death rate of the cell

Robust noise reduction by NC could impact various biological systems, particularly when small changes in molecular copy numbers significantly alter system output. To exemplify this, we utilized NC to reduce noise in the DNA repair system of *Escherichia coli (E. coli)* (Fig. 5a). In this system, when methyl methanesulfonate (MMS) enters the cell, it initiates DNA alkylation damage that can be repaired by the Ada protein. Ada is methylated when MMS enters the system, and methylated Ada induces expression of the *ada* gene, forming a positive feedback loop. Throughout this positive feedback, even a single Ada protein can trigger and amplify the expression of additional Ada. However, this happens only when at least one Ada molecule exists in the cell. For cells with no Ada, which is approximately 28% of the total population^19^, the positive feedback loop cannot be activated, leaving those cells unable to effectively respond to DNA damage (Fig. 5b). As a consequence, about one hour after MMS treatment, the *E. coli* population splits into two distinct groups depending on whether they succeed or fail to respond to DNA damage, resulting in a clearly bimodal distribution of Ada copy numbers (Fig. 5c). The group where the feedback loop is activated displays a cell death rate of 5% (Fig. 5c, d. blue), which is lower than the 15% cell death rate in the other group where the positive feedback loop fails to initiate (Fig. 5c, d. red).

**Figure 5.**
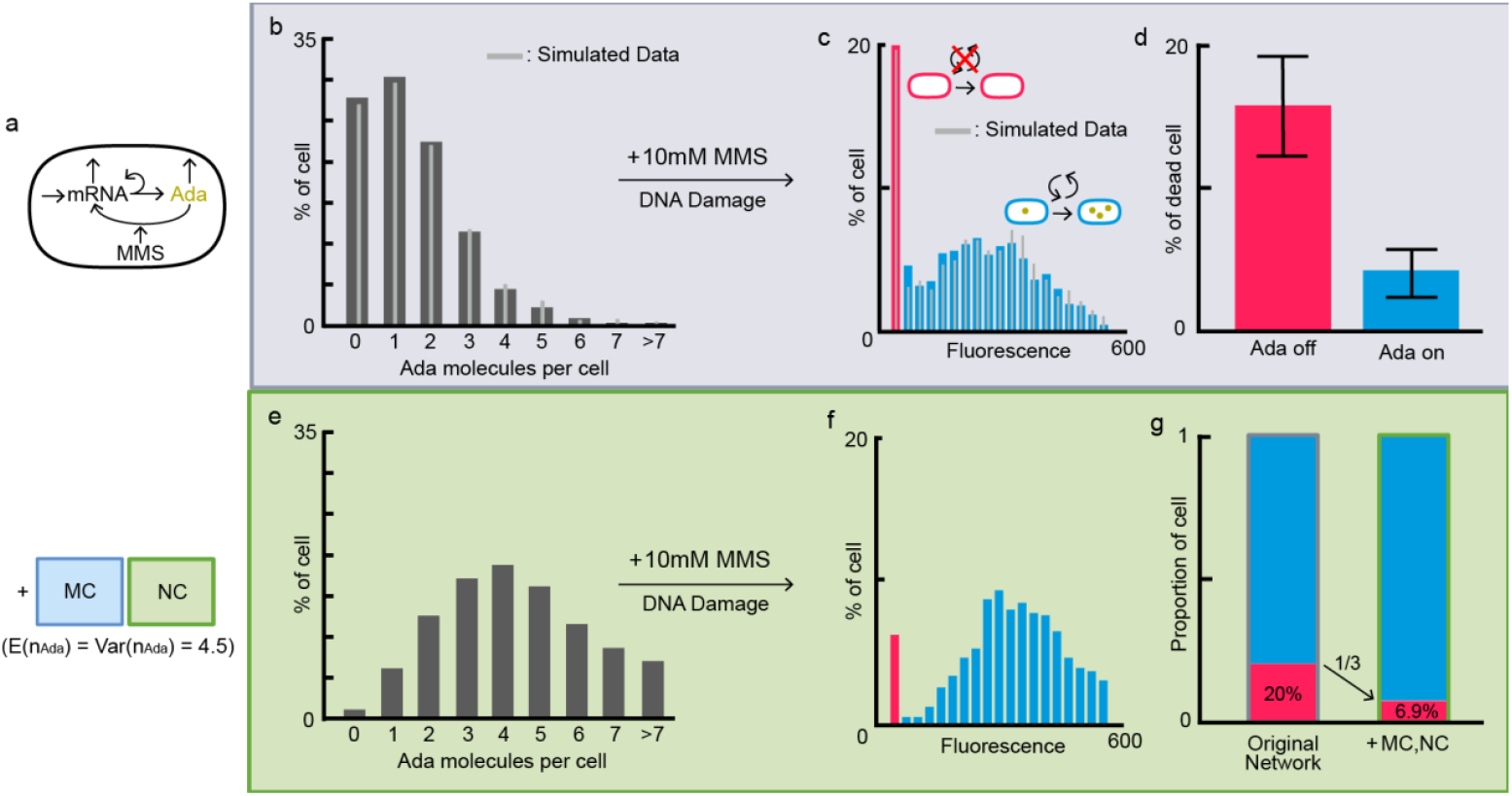
The noise controller reduces the death rate of a cell. **(a)** The DNA repair system of *E. coli*. An input of MMS activates a positive feedback loop only when the Ada protein is present. **(b)** The distribution of the number of Ada molecules per *E. coli* cell experimentally measured (gray bars) and reproduced from the model simulation (light gray lines). Simulating the reaction system shown in (a) successfully reproduced the experimental observation in Ref^19^. **(c)** The probability distribution of Ada-mYPet fluorescence per cell treated with MMS for 1 hour produced from the experiment (red and blue bars) and model simulation (gray lines). The probability distribution of Ada copy number has bimodality: the red area represents the cells without feedback response (Ada off), and the blue area represents the cells with feedback response (Ada on). **(d)** The ratio of dead cells after the MMS treatment measured in Ref^19^. Ada off cells show about four times higher death rate than Ada on cells. **(e)** The simulated probability distribution of the number of Ada molecules per cell with both the MC and NC, where both mean and variance were set at 4.5. Attaching both MC and NC reduces the probability of zero Ada molecules to nearly 1%. **(f)** The probability distribution of fluorescence per cell after the treatment with MMS. The red region indicates the area where Ada’s copy number is less than 900, the threshold which is obtained when reproducing the results in (c) using the same parameters. The probability of the red area decreased, and the blue area increased compared with (c). **(g)** The proportion of Ada on and Ada off cells before and after the control by MC and NC. The proportion of Ada off cells decreases when the MC and NC are attached. The experimental data shown in (b) - (d) are adopted from Ref^19^.

To investigate whether control by MC and NC could reduce the cell death rate by reducing the noise, we performed an *in silico* experiment to reproduce the experimental results. We first modeled a list of biochemical reactions to express a cell damage response process (Fig. 5a). The model was calibrated to fit the experimental data, and it successfully reproduced the distributions of Ada observed before and after MMS treatment (gray bars in Fig. 5b and 4c, see Methods for details). Notably, the model confirmed that the cells failing to respond to the damage accounted for approximately 20% of the population, which matches the experimental data (Fig. 5b).

With this calibrated model of a DNA repair system, we investigated whether combining MC and NC could reduce the occurrence of zero Ada levels. To do this, we first increased the mean copy number threefold using the MC. However, applying the MC alone led to an increase in noise, as expected, reducing the effectiveness of decreasing the probability of zero Ada molecules. To counteract this, we minimized the noise by setting the Fano factor to 1 using NC. The combined use of MC and NC dramatically reduced the probability of a cell having zero Ada—from 27% to 1% (a 27-fold reduction) (Fig. 5e). Consequently, the fraction of cells that failed to trigger an adequate DNA damage response was significantly reduced from 20% to approximately 7% (Fig. 5f, g). These results demonstrate that attaching MC and NC to the DNA repair system of *E. coli* could successfully reduce the death rate of the cell.

### MC and NC can be implemented in a biological circuit

We have confirmed that MC and NC can reduce the noise in real biological systems (Fig. 5), motivating their implementation in a real biological circuit. To this end, we conceptually designed a biological circuit of MC and NC inspired by previous studies that implemented MC in a biological circuit^6,20^. In the MC, the annihilation (or sequestration) reaction between two controller species (Fig. 6, part a) can be implemented using biological components such as sigma/anti-sigma factors, mRNA/sRNA pairs, scaffold/anti-scaffold interactions, or toxin/antitoxin systems^21^. Specifically, the RsiW/SigW protein pair was used in a previous study for *in vitro* realization via sequestration, wherein the two proteins bind to each other, thereby preventing their respective function^20^. Similarly, in the NC, the RseA/ECF pair could be employed to realize the sequestration of two controller species (Fig. 6, part b)^22^. However, the NC differs from the MC in two key structural aspects: the sensing reaction (Fig. 6, parts c-d) and the degradation-based actuation reaction (Fig. 6, part e). For the sensing reaction in NC, unlike in MC where the *X*_2_ monomer functions as a transcriptional activator (Fig. 6, part c), the *X*_2_ homodimer must take on this role (Fig. 6, part d). In other words, both the *X*_2_ monomer and the *X*_2_ homodimer should function as transcriptional activators for the MC and the NC, respectively. This can be realized using the transcription factor Pit-1^23^. The Pit-1 monomer binds to the prolactin (prl) enhancer of the *RsiW* gene (*Z*_2_), while the Pit-1 homodimer binds to the growth hormone (GH) enhancer of the *RseA* gene (*Z*_4_). Therefore, by fusing the Pit-1 DNA sequence with *X*_2_ and placing the prl DNA site next to *Z*_2_ (Fig. 6, part c) and the GH DNA site next to *Z*_4_ (Fig. 6, part d), this mechanism can be designed so that the *X*_2_ monomer activates the transcription of *Z*_2_, while the *X*_2_ dimer activates the transcription of *Z*_4_. Additionally, for the degradation-based actuation reaction, *Z*_4_ could be designed as a catalyst that induces the degradation of *X*_1_ (Fig. 6, part e). This could be realized by tagging *X*_1_ with a protein degradation tag (*pdt*), making it susceptible to degradation by the *Mesoplasma florum* (mf) Lon protease, thereby enabling tunable control of protein degradation^24^.

**Figure 6.**
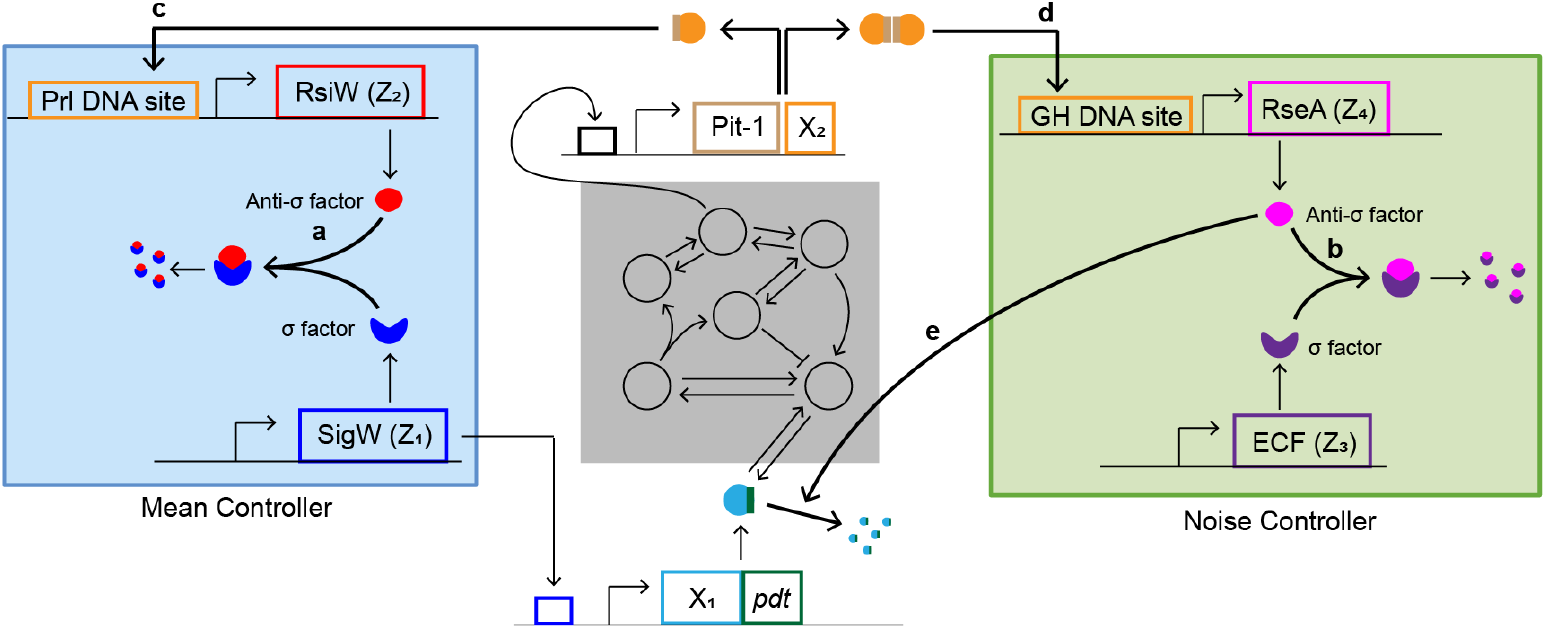
Schematic representation of a biological circuit implementing NC. (a) To biologically implement the MC (left, shaded blue box), we can use the sigma factor SigW (blue) as one controller species (*Z*_1_) and the anti-sigma factor RsiW (red) as another controller species (*Z*_2_), as done in the previous study^20^, because the expressed proteins (red circle and blue cup) sequester each other. (b) Similarly, to biologically implement the NC (right, shaded green box), we can use another sigma/anti-sigma factor such as ECF (purple) and RseA (pink) as controller species (*Z*_3_ and *Z*_4_, respectively). (c-d) To implement the sensing reactions of MC and NC (labelled c and d, respectively), we can fuse the DNA sequence of *X*_2_ with Pit-1 (light brown box, centre). For MC, the compound of Pit-1 (light brown rectangle; c) and monomer of *X*_2_ (orange circle; c) binds to the prolactin (Prl) enhancer site (left orange box; labelled c), thereby functioning as a transcription factor for the *Z*_2_ gene (red box; c). In this case, the reaction rate can be proportional to *X*_2_, as demonstrated in a previous study^20^. For NC, the compound of two Pit-1 (light brown rectangles; d) and a homodimer of *X*_2_ (orange circles; d) binds to the growth hormone (GH) DNA site, thereby functioning as a transcriptional factor for *Z*_4_ (pink box; d)^23^. In this case, the reaction rate can be a quadratic equation of *X*_2_, similar to MC (d). (e) To biologically implement the degradation-based actuation reaction of NC, we can tag *X*_1_ with a protein degradation tag (pdt). Once this construct is expressed, the tag induces *X*_1_ degradation by the *Mesoplasma florum* Lon (mf-lon) protease, which functions as *Z*_4_.

## Discussion

While single-cell level control requires RPA for the output noise level of the system, the module that could achieve this noise RPA had not been developed. To address this gap, we developed an NC to achieve noise RPA (Fig. 1d). Using this NC combined with the MC, we maintained both the output mean and noise levels even after a perturbation, achieving mean RPA and noise RPA simultaneously (Fig. 1e, f). Furthermore, by adjusting key parameters in the MC and NC, we reduced output noise up to a Fano factor of 1 (Fig. 2). This capability of the MC and NC applies as long as the combined system is ergodic with finite second moments, enabling noise reduction across various biological systems regardless of the sequestration and actuation parameters (Fig. 3). Notably, incorporating the MC and NC into a DNA repair system successfully reduced the failure rate of the DNA damage response (Fig. 5). Therefore, the NC enhances the precision and accuracy of biological controllers, enabling single-cell level control that the MC alone cannot achieve.

The MC and NC operate via distinct actuation mechanisms (Fig. S1). In particular, the MC relies on production-based actuation, producing an input species (X_1_ in Fig. S1) to regulate the output (X_2_ in Fig. S1). In contrast, the NC employs degradation-based actuation, directly degrading the input species. Importantly, production-based actuation alone cannot effectively control stochastic events in single cells. For example, if the MC maintains an average protein level of 10, stochastic fluctuation may result in cells reaching protein levels as high as 20. Reducing this level back to 10 is important for single-cell level control. To do this, the MC can only halt input production and wait for natural degradation. It is similar to a situation where lowering an object from a pulley requires the object itself to move downward (Fig. S1a). However, introducing an additional pulley, which pulls down the object, accelerates the process (Fig. S1b). This role of an additional pulley is analogous to the degradation-based actuation of the NC. Therefore, by integrating the NC, we can rapidly reduce excess protein levels (Fig. S1c), eventually reducing the number of cells whose protein levels exceed 10. Due to this property, we could restore the noise level from the increase caused by the MC.

Similar to the NC, several previous studies have proposed AIF-based controllers for noise reduction^14-16^ (Text S4). For example, output noise could be reduced by combining the MC and the negative feedback loop, where output species repress the production of input species^14^. Employing a similar negative feedback loop as an alternative to the MC’s actuation achieved noise reduction below the original level^15,16^. Some of these achieved a Fano factor below 1, but after parameter perturbations, they typically failed to maintain low noise levels (Text S4, Fig. S6). To resolve this limitation, we developed the NC, which senses the output levels via the dimerization of output species (Fig. 1d). This approach ensures the control of the second moment of the output (See Methods for more details), which leads to the noise RPA (Fig. S6b, d, f). In light of this, the dimerization of output species is likely a key feature for achieving noise RPA. Identifying natural systems that exhibit noise RPA and investigating whether they share this dimerization feature presents a promising direction for future research.

In natural systems, noise typically scales with the square root of the mean, a principle referred to as the 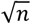 law from Schrödinger^25^. In line with this principle, systems with a Fano factor of 1 have been extensively studied^26,27^. However, many biological systems exhibit Fano factors greater than 1. For example, simple gene expression models theoretically exhibit a Fano factor greater than 1^28^. Furthermore, empirical analysis of 1081 different proteins in *E. coli* confirmed that most proteins have a Fano factor greater than 1^29^. Thus, a Fano factor of 1 may represent the lower limit of noise in natural systems, meaning that NC is the module capable of reducing noise to this fundamental limit of natural systems. Nevertheless, a deeper theoretical investigation of the Fano factor = 1 limit is warranted. A lower bound on the noise of an output species regulated by a feedback loop has been established^30^, by using the theory of information transmission. Applying this theory to our scenario, where an output species is regulated by MC and NC, remains an important direction for future work.

While the Fano factor of 1 may represent the natural lower bound of noise, recent studies have reported cases where the Fano factor drops below 1. For instance, the telegraph model of gene expression systems can achieve Fano factors below 1 if transition rates between on/off gene states are fast and a negative feedback loop is present^31^. In addition, another study proved that any discrete probability distribution, including those with Fano factors below 1, can be approximated using the stationary distribution of a chemical reaction network^32^. Therefore, by applying these studies, the Fano factor can be reduced below 1. Thus, investigating the feasibility of this remains a promising direction for future work.

Despite the advantages of MC and NC, their species—particularly *Z*_1_ and *Z*_4_ —can accumulate indefinitely when the target Fano factor is below 1 (Fig. 2d). Such accumulation of controller species can be detrimental to cells and may impair the proper functioning of MC and NC. A promising strategy to prevent this issue is the anti-windup approach proposed by Filo et al.^33^, which has been shown to effectively prevent the accumulation of MC species. Using this anti-windup topology, we confirmed that the accumulation of *Z*_1_ and *Z*_4_ can also be effectively prevented (see Text S1 and Fig. S2 for details). However, incorporating anti-windup led to a deviation between the actual and target Fano factors (Fig. S2), as it alters the Fano factor formula described in Fig. 2a (Text S1). As the altered Fano factor formula may depend on the parameters of original networks, it remains uncertain whether noise RPA would still hold when anti-windup is applied. Therefore, developing a method that prevents controller species accumulation via anti-windup while also theoretically ensuring the noise RPA of MC and NC represents a promising direction for future work.

Through numerical simulations of several open-loop networks with various parameter values (Fig. 2, 3), we observed that the reaction network associated with NC is stable when the target Fano factor exceeds 1, and unstable when the target Fano factor is below 1. To establish this claim rigorously, we proved that in some parameter regions where the target Fano factor is less than 1, the associated continuous-time Markov chain is transient (Text. S3).

Furthermore, we found that showing sufficient conditions for ergodicity of the controlled system may lead to new theoretical approaches. One clear difficulty comes from the production-based control by *Z*_1_ → *Z*_1_ +*X*_1_ in MC and the degradation-based control by *Z*_4_ +*X*_1_ → *Z*_4_ in NC. As they give opposite drifts, one may think of using a piecewise Lyapunov function as shown in previous study^6^. However, due to the non-linearity of the degradation reaction in NC, it is challenging to control the drifts around the boundaries between the regions where a piecewise Lyapunov function is defined differently. Another common approach is to use a skeleton chain, which is a discrete-time Markov chain obtained by sampling the original Markov chain at certain time points^35^. This approach is often employed to exploit negative drifts that are not present initially but emerge after some time. However, this method does not appear to be easily applicable here. For example, when the positive drift induced by 2*X*_2_ → 2*X*_2_ + *Z*_4_ dominates, this dominance is expected to last for *O*(log *n*) time if the initial amount of *X*_2_ is *n*. It would be highly challenging to balance the positive drift accumulated over an *O*(log *n*) time interval with a dominant negative drift that appears only afterward. Consequently, our analysis indicates that proving the ergodicity of the NC requires a fundamentally new analytical framework rather than conventional techniques typically used for reaction networks. Future work toward proving the ergodicity and finiteness of the moment of a reaction network with NC may create a new theoretical foundation for future studies in stochastic reaction networks.

One of the key challenges in implementing the NC as a biological circuit is the saturation effect. Although the sensing reaction follows mass-action kinetics in this study, biochemical reaction rates often follow a Hill-function form in reality. This issue can be addressed by tuning the dissociation constant of the sensing reaction (Text S5, Fig. S7). In the supplementary text, we checked that the NC with saturation effect could achieve the noise RPA and set-point tracking by larger dissociation constant (Fig. S7). To realize a large dissociation constant in practice, one could modify the base sequence of the GH DNA site (Fig. 6d) or engineer the DNA-binding domain of the *X*_2_ homodimer (Fig. 6d) to make the binding affinity low.

While the proposed biological circuits for MC and NC appear feasible in principle (Fig. 6), their practical implementation poses several challenges. One issue is the incompatibility between cellular systems: while Pit-1 and its promoter have been studied in eukaryotic cells (Fig. 6c,d)^23^, the other components of the circuit are based on prokaryotic systems^6,20^. This incompatibility may interrupt the proper function of Pit-1, which is key for implementing the sensing reaction of the MC and NC (Fig. 6, c,d). To address this, a prokaryotic alternative with similar regulatory function must be identified. Another major bottleneck is identifying suitable candidates for *Z*_4_ (Fig. 6b,e). In particular, *Z*_4_ must both promote the degradation of the input species, *X*_1_ (Fig. 6e) and also undergo sequestration with *Z*_3_ (Fig. 6b). Identifying a single protein that fulfills both of these functions is challenging. As a potential solution, we can engineer a chimeric protein by combining the DNA sequence of RseA, which detects the ECF (Fig. 6b), and the DNA sequence of mf-Lon protease, which facilitates the degradation (Fig. 6e)^36^. As another alternative, an RNA-based expression control system could also be implemented^37^. For example, *X*_1_, *Z*_3_ and *Z*_4_ could be replaced with engineered small RNAs. In this design, *Z*_4_ functions as an aptazyme-containing antisense RNA: its aptazyme domain cleaves the *X*_1_ transcript, degrading the *X*_1_ (Fig. 6e), while its antisense RNA region enables mutual hybridization with a complementary small RNA, *Z*_3_, resulting in their annihilation (Fig. 6b), as proposed previously^34^. However, these approaches require further validation to confirm that the engineered chimeric protein or aptamer-based circuit can perform both desired functions properly.

Although these challenges have hindered the experimental validation of NC, future studies confirming the NC’s biological feasibility are essential. Implementing NC biologically and demonstrating single-cell level control would establish NC’s practical applicability. If successful, the MC and NC could be widely used in various areas requiring precise single-cell control, such as cancer therapy and tuberculosis treatment. In addition to the biological feasibility of the NC, its theoretical feasibility also warrants further investigation. For example, ergodicity is required for noise RPA, yet the precise conditions of the open-loop structure and the parameters for ergodicity, including the admissible perturbation size, are not fully understood. The ergodicity condition has been analytically expressed for the system that only used MC^6^, and extending this method to the system combined with both MC and NC will provide further important insights (We provide a proof of transience for the system with MC and NC under a restricted parameter set—a subset of the regime in which the target Fano factor is < 1—in Text S3.).

## Methods

### Derivation of the stationary mean and variance of the network combined with MC and NC

To regulate the second moment of the system output as well as its first moment, we developed the NC inspired by the MC, as follows:

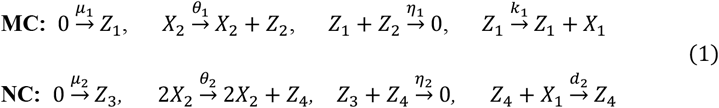

where *X*_1_ and *X*_2_ are the input and output species of the target system, respectively. Then, the dynamics of the first-order moment of *Z*_1_ – *Z*_4_ could be expressed as an ordinary differential equation derived from the chemical master equation.

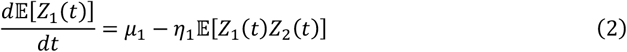

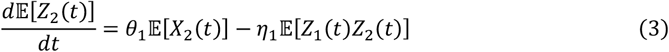

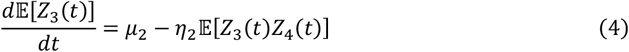

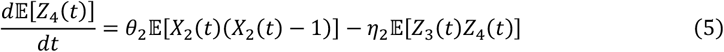

where *X*_1_(*t*),*X*_2_(*t*), *Z*_1_(*t*), *Z*_2_(*t*), *Z*_3_(*t*) and *Z*_4_(*t*) denotes the molecular counts of the *X*_1_,*X*_2_, *Z*_1_, *Z*_2_, *Z*_3_ and *Z*_4_. Then, by subtracting (3) from (2) and (5) from (4), we could derive the new equations:

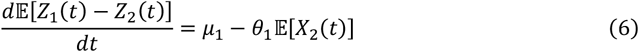

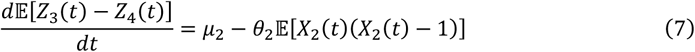

If we assume the *ergodicity* of a system, where *ergodicity* of the system implies that the probability distribution of *X*_1_,*X*_2_, *Z*_1_, *Z*_2_, *Z*_3_ and *Z*_4_ converge to a stationary distribution as *t* → ∞. This leads to convergence of the first and the second moments of *Z*_1_, *Z*_2_, *Z*_3_ and *Z*_4_ as well if they are finite. When their first moment reaches a steady-state, the left-hand side of equations (6) and (7) becomes zero. As a result, we get the following equations for the first and second moment of *X*_2_:

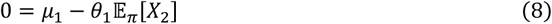

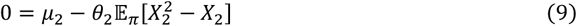

where π is the stationary distribution of the system. Then, we can derive the formula of stationary mean of *X*_2_ as 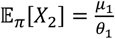 from (8), and the formula of stationary variance of *X*_2_ as Var 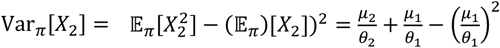 from (9). Therefore, we get the following formula of Fano factor of *X*_2_ at the stationary distribution:

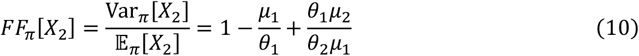

### Simulation details for the heatmaps

We assessed the ability of the MC and the NC to reduce noise to a Fano factor of 1 by generating heatmaps depicting the ratio between the simulated and target Fano factors in the original network combined with the MC and the NC (Figs. 2 and 3). To do this, we performed stochastic simulations for various target mean and Fano factor values. The target mean ranged from 2 to 5 in increments of 0.2, while the target Fano factor varied from 0.1 to 2.5 in increments of 0.1, except in Figs. 3b and 3d. Specifically, in Fig. 3b, when *k*_1_ = 10, the target Fano factor ranged from 0.3 to 7.5 with a step size of 0.3, whereas in Fig. 3d, it ranged from 0.75 to 1.5 with a step size of 0.03.

For each target mean and Fano factor pair, we simulated 1000 trajectories of *X*_2_ in the combined system (Fig. 1a), initializing all species at zero and running each simulation for 10,000 seconds. The Fano factor of 1000 trajectories was computed, forming a time series of Fano factor values spanning 10,000 seconds. The last 2,000 seconds of this time series was averaged to obtain the converged Fano factor value. Finally, this value was divided by the target Fano factor and visualized as heatmaps (Figs. 2 and 3).

In Fig. 3, we utilized two different original networks: original network 2, which represents a dimerization process (Fig. 3c), and original network 3, which includes a bimolecular reaction process (Fig. 3d). The original network 2 consisted of the following reactions:

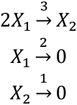

where *X*_1_ is the input species and *X*_2_ is the output species. The original network 3 consisted of the following reactions:

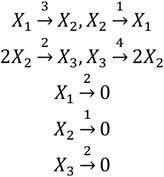

where *X*_1_ is both input and output species, as done in a previous study^20^.

### Details of bimodality analysis using telegraph and toggle switch models

We demonstrated the MC and NC’s ability to remove the bimodality using the telegraph model (Fig. 4a-c) and toggle switch model (Fig. 4d-f), which exhibits the bimodal output stationary distribution from fast and slow production modes. Here, the telegraph model consists of the following reactions:

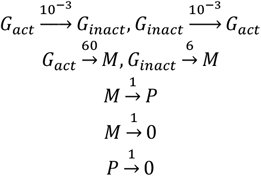

where *G*_act_, *G*_inact_, *M*, and *P* denote the active gene, inactive gene, mRNA, and protein, respectively. The output species was *P*, while the input species was *M*. The toggle switch model consists of the following reaction:

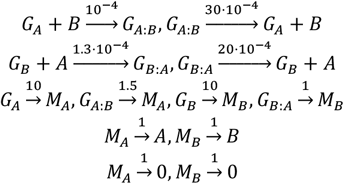

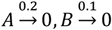

where *G*_A_, *G*_A:B_, *G*_B_, *G*_B:A_, *M*_A_, *M*_B_, *A*, and *B* denote the gene A, gene A bind with B, gene B, gene B bind with A, mRNA of A, mRNA of B, protein A, and protein B, respectively. The output species was *A*, while the input species was *M*_A_.

For the telegraph model, we simulated 1,000 trajectories of output species (i.e., *P*) in the telegraph model (Fig. 4b (i)), the telegraph model with MC (Fig. 4b (ii)), and the telegraph model with MC and NC (Fig. 4b (iii)), initializing all species at zero except one active gene and running each simulation for 30,000 seconds. The output stationary distribution was obtained from the final state of *P* in each trajectory (value at 30,000 s). To define the fast-mode production propensity (Fig. 4c), at each time point *t*, we computed the mean protein level across those trajectories in which the gene was active at *t*; for example, the fast-mode propensity at *t* = 1,000*s* is the average of *P* over all trajectories with the gene active at 1,000s. Applying the same method to the cases where the gene was inactive, we obtained the propensity in the slow production mode.

For the toggle switch model, we simulated 1,000 trajectories of output species (i.e., *A*) in the toggle switch model (Fig. 4e (i)), the toggle switch model with MC (Fig. 4e (ii)), and the toggle switch model with MC and NC (Fig. 4e (iii)), initializing all species at zero except one unbound gene A and one unbound gene B and running each simulation for 20,000 seconds. The output stationary distribution was obtained from the final state of *A* in each trajectory (value at 20,000 s). To define the fast-mode production propensity (Fig. 4f), at each time point *t*, we computed the mean level of *A* across those trajectories in which the gene was not bound with *B* at *t*; for example, the fast-mode propensity at *t* = 1,000*s* is the average of *B* over all trajectories with the gene not bound with *B* at 1,000s. Applying the same method to the cases where the gene was bound with *B*, we obtained the propensity in the slow production mode.

### Details of the *in silico* experiment with the DNA repair system

For the *in silico* experiment in Fig. 5, we constructed a DNA repair system with the following biochemical reactions.

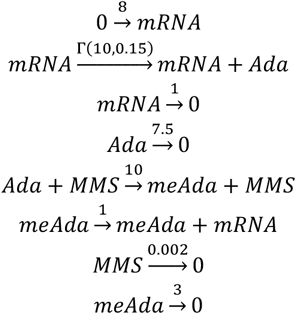

The parameters of each reaction were selected to reproduce the experimental results reported in Uphoff et al.^19^ (Fig. 5b,c). For the parameter of the Ada translation reaction (the second reaction in the above diagram), we introduced a gamma distribution to account for the heterogeneity among the cells, as done in the previous study^38^. In addition, the propensity of the methylation of Ada was expressed using the hill function, 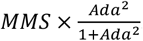. For this original network of a DNA repair system, we combined the MC and NC composed of the following reactions:

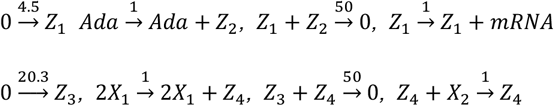

The reaction parameters of MC and NC were chosen so that the target mean is three times bigger than the original value (from 1.5 to 4.5) and the target Fano factor is close to 1 (1.01). The DNA repair system combined with MC and NC was simulated until it reached the stationary distribution, and then 100 copies of MMS entered the system. Simultaneously, MC and NC stopped their operation (i.e. all the parameters in MC and NC dropped to 0 immediately after the MMS treatment) to ensure the amplification of the Ada copy number through positive feedback.

To scale the copy number of Ada to fluorescence, we matched the thresholds that distinguish Ada on/off cells. In Uphoff et al., Ada-off cells comprised 20% of the population, corresponding to one bar in the fluorescence histogram shown in Fig. 1c of the reference^19^. Because the six bars in that figure covered a fluorescence range of 200-400, the width of one bar can be estimated as 100/3. This implies that the threshold distinguishing Ada-on/off cells is 100/3 in fluorescence units. On the other hand, our simulations defined the threshold as an Ada copy number of fewer than 900 molecules (Fig. 5f legend). Accordingly, we set 900 Ada molecules to be equivalent to a fluorescence value of 100/3. The scaled fluorescence distribution after 5 seconds from the MMS treatment was captured and visualized in Fig. 5f.

## Data and Code Availability

The codes underlying this work will be made publicly available upon acceptance.

## Acknowledgements

The research was supported by the following agencies and institutions: the Institute for Basic Science (grant no. IBS-R029-C3, J.K.K.); Samsung Science and Technology Foundation (grant no. SSTF-BA1902-01, J.K.K.); the National Research Foundation of Korea (NRF) grant funded by the Korea government (MSIT) (Grant Nos. 2022R1C1C1008491 and RS-2023-00219980, J.K.); the NRF grant funded by MSIT (Grant No. 2021R1A6A1A10042944, J.K.); POSCO HOLDINGS research fund (Grant No. 2022Q019, J.K.); Samsung Electronics Co., Ltd. (Grant No. IO230407-05812-01, J.K.); Korea Health Technology R&D Project through the Korea Health Industry Development Institute (KHIDI), funded by the Ministry of Health & Welfare, Republic of Korea (grant number : RS-2024-00406488, J.K.); the NRF grant funded by MSIT (RS-2021-NR056566, B.-K.C.). We thank Helen Pickersgill in the Life Science Editors Foundation for assistance in English language improvement.

## Author Contributions Statement

Simulation: D.L., S.M.

Analysis of results: All authors

First draft: D.L., S.M., J.K., J.K.K.

Writing: All authors

## Competing Interest Statements

The authors declare no competing interests.

## Notes

### Competing Interest Statement

The authors have declared no competing interest.

## References

1 Arcus, V. L. et al. On the temperature dependence of enzyme-catalyzed rates. Biochemistry 55, 1681–1688 (2016). 10.1021/acs.biochem.5b01094

2 Khammash, M. H. Perfect adaptation in biology. Cell Syst 12, 509–521 (2021). 10.1016/j.cels.2021.05.020

3 Araujo, R. P. & Liotta, L. A. The topological requirements for robust perfect adaptation in networks of any size. Nat Commun 9, 1757 (2018). 10.1038/s41467-018-04151-6

4 Ferrell, J. E., Jr. Perfect and near-perfect adaptation in cell signaling. Cell Syst 2, 62–67 (2016). 10.1016/j.cels.2016.02.006

5 Ma, W., Trusina, A., El-Samad, H., Lim, W. A. & Tang, C. Defining network topologies that can achieve biochemical adaptation. Cell 138, 760–773 (2009). 10.1016/j.cell.2009.06.013

6 Briat, C., Gupta, A. & Khammash, M. Antithetic integral feedback ensures robust perfect adaptation in noisy biomolecular networks. Cell Syst 2, 15–26 (2016). 10.1016/j.cels.2016.01.004

7 Gupta, A. & Khammash, M. Universal structural requirements for maximal robust perfect adaptation in biomolecular networks. Proc Natl Acad Sci U S A 119, e2207802119 (2022). 10.1073/pnas.2207802119

8 Yi, T. M., Huang, Y., Simon, M. I. & Doyle, J. Robust perfect adaptation in bacterial chemotaxis through integral feedback control. Proc Natl Acad Sci U S A 97, 4649–4653 (2000). 10.1073/pnas.97.9.4649

9 Qian, Y. & Del Vecchio, D. Realizing ‘integral control’ in living cells: how to overcome leaky integration due to dilution? J R Soc Interface 15 (2018). 10.1098/rsif.2017.0902

10 Agrawal, D. K., Marshall, R., Noireaux, V. & Sontag, E. D. In vitro implementation of robust gene regulation in a synthetic biomolecular integral controller. Nat Commun 10, 5760 (2019). 10.1038/s41467-019-13626-z

11 Skeel, R. T. & Khleif, S. N. Handbook of cancer chemotherapy. (Lippincott Williams & Wilkins, 2011).

12 Gomez, J. E. & McKinney, J. D. M. tuberculosis persistence, latency, and drug tolerance. Tuberculosis (Edinb) 84, 29–44 (2004). 10.1016/j.tube.2003.08.003

13 Olsman, N. & Paulsson, J. Universal control in biochemical circuits. NATURE 570 (2019).

14 Briat, C., Gupta, A. & Khammash, M. Antithetic proportional-integral feedback for reduced variance and improved control performance of stochastic reaction networks. J R Soc Interface 15 (2018). 10.1098/rsif.2018.0079

15 Kell, B., Ripsman, R. & Hilfinger, A. Noise properties of adaptation-conferring biochemical control modules. Proc Natl Acad Sci U S A 120, e2302016120 (2023). 10.1073/pnas.2302016120

16 Filo, M., Kumar, S. & Khammash, M. A hierarchy of biomolecular proportional-integral-derivative feedback controllers for robust perfect adaptation and dynamic performance. Nat Commun 13, 2119 (2022). 10.1038/s41467-022-29640-7

17 Munsky, B., Neuert, G. & van Oudenaarden, A. Using gene expression noise to understand gene regulation. Science 336, 183–187 (2012). 10.1126/science.1216379

18 Hong, H., Kim, J., Ali Al-Radhawi, M., Sontag, E. D. & Kim, J. K. Derivation of stationary distributions of biochemical reaction networks via structure transformation. Commun Biol 4, 620 (2021). 10.1038/s42003-021-02117-x

19 Uphoff, S. et al. Stochastic activation of a DNA damage response causes cell-to-cell mutation rate variation. Science 351, 1094–1097 (2016). 10.1126/science.aac9786

20 Aoki, S. K. et al. A universal biomolecular integral feedback controller for robust perfect adaptation. Nature 570, 533–537 (2019). 10.1038/s41586-019-1321-1

21 Gupta, A. & Khammash, M. in IEEE Conf. Decis. Control (CDC). 2808–2813 (IEEE).

22 Campbell, E. A. et al. Crystal structure of Escherichia coli sigmaE with the cytoplasmic domain of its anti-sigma RseA. Mol Cell 11, 1067–1078 (2003). 10.1016/s1097-2765(03)00148-5

23 Holloway, J. M., Szeto, D. P., Scully, K. M., Glass, C. K. & Rosenfeld, M. G. Pit-1 binding to specific DNA sites as a monomer or dimer determines gene-specific use of a tyrosine-dependent synergy domain. Genes Dev 9, 1992–2006 (1995).

24 Cameron, D. E. & Collins, J. J. Tunable protein degradation in bacteria. Nat Biotechnol 32, 1276–1281 (2014). 10.1038/nbt.3053

25 Schrodinger, E. What is life. (Cambridge University Press, 2012).

26 Anderson, D. F., Craciun, G. & Kurtz, T. G. Product-form stationary distributions for deficiency zero chemical reaction networks. Bull Math Biol 72, 1947–1970 (2010). 10.1007/s11538-010-9517-4

27 Whittle, P. Systems in stochastic equilibrium. (John Wiley & Sons, Inc., 1986).

28 Hilfinger, A. & Paulsson, J. Separating intrinsic from extrinsic fluctuations in dynamic biological systems Proc Natl Acad Sci U S A 108, 12167–12172 (2011). 10.1073/pnas.1018832108

29 Taniguchi, Y. et al. Quantifying E. coli proteome and transcriptome with single-molecule sensitivity in single cells. Science 329, 533–538 (2010). 10.1126/science.1188308

30 Lestas, I., Vinnicombe, G. & Paulsson, J. Fundamental limits on the suppression of molecular fluctuations. Nature 467, 174–178 (2010). 10.1038/nature09333

31 Ramos, A. F., Hornos, J. E. & Reinitz, J. Gene regulation and noise reduction by coupling of stochastic processes. Phys Rev E Stat Nonlin Soft Matter Phys 91, 020701 (2015). 10.1103/PhysRevE.91.020701

32 Cappelletti, D., Ortiz-Muñoz, A., Anderson, D. F. & Winfree, E. Stochastic chemical reaction networks for robustly approximating arbitrary probability distributions. Theor Comput Sci, 64–95 (2020).

33 Filo, M., Gupta, A. & Khammash, M. Anti-windup strategies for biomolecular control systems facilitated by model reduction theory for sequestration networks. Sci Adv 10, eadl5439 (2024). 10.1126/sciadv.adl5439

34 Agrawal, D. K. et al. Mathematical modeling of RNA-based architectures for closed loop control of gene expression. ACS Synth Biol 7, 1219–1228 (2018). 10.1021/acssynbio.8b00040

35 Anderson, D. F., Cappelletti, D. & Kim, J. Stochastically modeled weakly reversible reaction networks with a single linkage class. Journal of Applied Probability 57, 792–810 (2020).

36 Campbell, R. K., Bergert, E. R., Wang, Y., Morris, J. C. & Moyle, W. R. Chimeric proteins can exceed the sum of their parts: implications for evolution and protein design. Nat Biotechnol 15, 439–443 (1997). 10.1038/nbt0597-439

37 Zhou, S. et al. Trans-acting aptazyme for conditional gene knockdown in eukaryotic cells. Mol Ther Nucleic Acids 33, 367–375 (2023). 10.1016/j.omtn.2023.07.014

38 Cortez, M. J., Hong, H., Choi, B., Kim, J. K. & Josic, K. Hierarchical Bayesian models of transcriptional and translational regulation processes with delays. Bioinformatics 38, 187–195 (2021). 10.1093/bioinformatics/btab618

